# A single dose SARS-CoV-2 simulating particle vaccine induces potent neutralizing activities

**DOI:** 10.1101/2020.05.14.093054

**Authors:** Di Yin, Sikai Ling, Xiaolong Tian, Yang Li, Zhijue Xu, Hewei Jiang, Xue Zhang, Xiaoyuan Wang, Yi Shi, Yan Zhang, Lintai Da, Sheng-ce Tao, Quanjun Wang, Jianjiang Xu, Tianlei Ying, Jiaxu Hong, Yujia Cai

## Abstract

Coronavirus disease 2019 (COVID-19) is caused by severe acute respiratory syndrome coronavirus 2 (SARS-CoV-2) for which a vaccine is urgently needed to control its spreading. To facilitate the representation of a native-like immunogen without being infectious, here, we reported a SARS-CoV-2 vaccine candidate (designated ShaCoVacc) by incorporating spike-encoding mRNA inside and decorating spike protein on the surface of the virus simulating particles (VSPs) derived from lentiviral particles. We characterized the mRNA copy number, glycosylation status, transduction efficiency, and innate immune property of the new vaccine platform. Importantly, we showed the ShaCoVacc induced strong spike-specific humoral immune responses and potent neutralizing activities by a single injection. Additionally, we disclosed the epitopes of spike-specific antibodies using peptide microarray and revealed epitopes susceptible to specific neutralizing antibodies. These results support further development of ShaCoVacc as a candidate vaccine for COVID-19 and VSP may serve as a new vaccine platform for emerging infectious diseases.

---

The coronavirus disease-19 (COVID-19) has been rapidly becoming a globe pandemic since its first report in Wuhan, China, in the December of 2019^1^. The disease is caused by Severe Acute Respiratory Syndrome Coronavirus 2 (SARS-CoV-2) which was effectively transmitted from human to human, with influenza-like symptoms ranging from mild disease to severe lung injury and multi-organ failure, eventually causing death, especially in aged patients with co-morbidities^2,3^. Vaccines are considered to be one of the most effective ways to terminate the pandemic and help to restore the global economy^4,5^. By May 12, at least six SARS-CoV-2 vaccine candidates have entered the clinical stage basing on different platforms, i.e. mRNA, adenovirus, lentivirus, and plasmid DNA^6^. However, published information regarding the neutralizing efficacy of each vaccine is still very limited except an inactivated vaccine for which, however, multiple vaccinations are needed to get high EC50 titer^7,8^.

In the past decades, numerous vaccine platforms have been approved for markets or clinical trials. Live attenuated vaccines are weakened pathogens that induce strong humoral and cellular immune response but also with infection risks especially for immune-compromised people^9,10^. Inactivated vaccines are killed pathogens with intact structure and destroyed genetic materials – therefore, less risky, but also less efficacy^9,10^. Protein subunit vaccine, viral vector vaccine, and nucleic acid vaccine (DNA and mRNA) are generally safe, but difficult to reflect the conformational structure of viral immunogen in its nature^11,12^. Virus-like particles (VLPs) are hulk particles with the unique ability to present viral spikes in their natural conformation and elicit conformation-dependent neutralizing antibodies^13–16^. A closer mimicking of a pathogen would be a VLP with spikes on its surface and antigen-encoding nucleic acids inside. It is still unknown which vaccine platform will actually work for SARS-CoV-2, making the development of new vaccine platforms of great importance. As neutralizing antibodies have been detected in convalescents of COVID-19^17–20^, a SARS-CoV-2 simulating vaccine may present the antigens to the immune systems in much the same way that it would be presented by an authentic virus, thereby provoking a similar effective immune response.

In this study, we designed a candidate vaccine by encapsulating spike within and decorated onto the surface of virus simulating particles (VSPs) derived from lentiviral particles in the form of mRNA and protein, respectively, simulating the wild-type SARS-CoV-2. We hypothesized this could be achieved by co-expression of a full-length spike during the production of mRNA-carrying lentiviral particle which we have recently developed for time-restricted gene editing (Manuscript under revision). To package the full-length spike mRNA into VSPs, we designed a spike construct expressing the spike protein with 6X MS2 stem loop repeats on its transcripts which would allow the spike mRNA to be packaged into VSPs via interaction with MS2 coat fused GagPol (Fig. 1a). Meanwhile, as an envelope protein, the spike protein would automatically assemble into the membrane of virus simulating particles (Fig. 1b).

**Figure 1.**
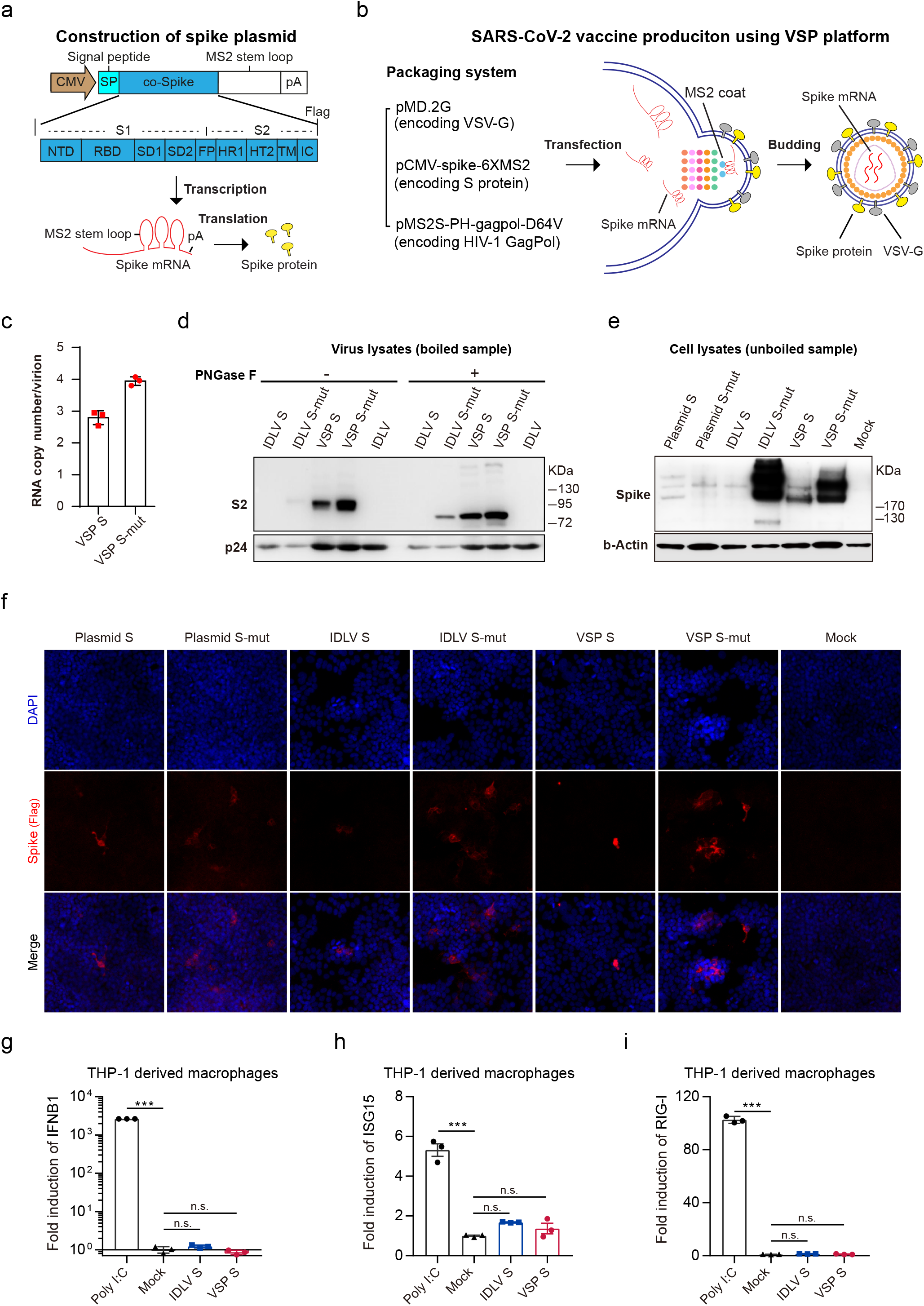
Construct and characterization of a SARS-Cov-2 simulating particle vaccine. **a**, Construction of mRNA-encoding plasmid which transcribes a MS2 stem loop-containing spike mRNA. The spike mRNA and protein will be packaging into lentiviral particles via the RNA-coat protein interaction and self-assembly, respectively. NTD, N-terminal domain; RBD, receptor binding domain; SD1 and SD2, subdomain 1 and 2; FP, fusion peptide; HR1 and HR2, heptad repeat 1 and 2; TM, transmembrane domain; CT, cytoplasmic tail. **b**, Illustration of the production process of the SARS-CoV-2 vaccine using VSP platform. **c**, Copy number of spike mRNA in each VSP particle. The copy number was detected by absolute quantification RT-qPCR and normalized to IDLV-spike-mut (2 copies RNA per virion). **d**, Western blot analysis of the spike protein in the virion treated with/without PNGase F. IDLV use as a control. 100 ng p24 for each vector. **e**, Western blot analysis of the spike protein expression. 293T cells were collected 36 hr after transfection or transduction. 300 ng p24 virus for each well. **f**. Confocal analysis of spike protein expression. 293T cells were fixed 36 hr after transfection or transduction. **g-i**, Innate immune response induced by VSP in THP-1 derived macrophages. Cells were harvested for IFNB1, ISG15 and RIG-I analysis by RT-qPCR 6 hr after transduction 150 ng p24 per well for each virus and 1.5 μg poly I:C per well as positive controls. S represents spike. One-way ANOVA with Dunnett’s *post hoc* tests were performed, ***P< 0.001, n.s.=non-significant.

To make sure high transcription and translation in the producer cells, we used a CMV promoter to drive spike expression (Fig. 1a). In addition, the spike sequence was codon-optimized and equipped with a signal peptide from the human heavy chain of IgE to boost translation and immunogenicity (Fig. 1a)^4^. We also introduced two proline substation mutations in the S2 of spike sequence which have been reported to increase the expression of spike and enhance the production of neutralizing antibodies (Fig. 1a)^21,22^. The resulting spike transcript contains a MS2 stem loop between the stop codon and the polyA sequence so that it could be packaged into lentiviral particles via direct interaction with its cognate MS2 coat protein localized in the N-terminus of GagPol polyprotein. As VSV-G coated lentiviruses are efficiently taken up by antigen-presenting cells and show the high immunogenicity of antigens^23^, we also included the VSV-G encoding plasmid (pMD.2G) in the production process so that the surface of resulting VSPs will be covered by both spike and VSV-G (Fig. 1b).

To examine if spike mRNA has been packaged into lentiviral particle as designed, we performed RT-qPCR and found 3 or 4 copies of spike mRNA on average for each VSP (Fig. 1c). To verify whether spike proteins have been assembled into VSPs and their glycosylation status, we performed Western blot on the lysates of VSP with integration-defective lentiviruses (IDLVs) as controls (Fig. 1d). We found successful decoration of spikes both with and without mutations on the VSPs while more mutant spikes could be loaded. As glycosylation impacts the immunogenicity and immunodominance of a vaccine^24^, we examined the glycosylation status of the spike on the surface of VSPs. Notably, the S2 bands shifted downwards after PNGase F treatment indicating that the spikes on VSPs were modified by N-linked glycosylations consistent with the resent finding for SARS-CoV-2 revealed by mass spectrometric approach (Fig. 1d)^25^. Next, we transduced VSPs to 293T cells and evaluated the expression of spikes. We harvested the cells 36 hr postinfection without boiling the samples for Western blot to avoid potential aggregation of the spike protein. 293T cells were not infected by SARS-CoV-2 unless supplemented with hACE2^26^. Here, we still observed the expression of spikes in 293T cells from VSPs suggesting that VSV-G were co-assembled into VSPs, thereby, broadened their tropism (Fig. 1e). We found two major bands for spikes which were likely glycosylated full-length singlet spike and its dimeric/trimeric forms (Fig. 1e). Additionally, we confirmed the expression of spike using confocal analysis of transfected or transduced 293T cells (Fig. 1f). To examine the innate immune property of VSP, we used THP-1 derived macrophages as a model of nucleic sensing and found no significant increase of the type I interferon (IFN) and IFN-stimulated genes ISG-15, and retinoic acid-inducible gene I (RIG-I) (Fig. 1g-i). As VSP-spike-mut incorporates both mRNA and protein of spike more efficiently than the wild-type counterpart, we therefore chose it as a vaccine candidate (designated ShaCoVacc) for in vivo evaluation.

To access the immunogenicity of ShaCoVacc, we injected the vaccine candidate to C57BL/6J mice (n=6 for each group) via footpad (Fig. 2a). We performed enzyme-linked immunosorbent assay (ELISA) using the sera from mice two weeks after vaccination to get access to the spike-specific IgG. As shown in Fig. 2b, we observed significant elicitation of the spike-specific IgG. To evaluate the production of neutralizing antibodies, we performed the neutralizing assay using spike pseudotyped HIV which encodes firefly luciferase - a well-established pseudovirus neutralization assay^27^. Interestingly, a single injection of ShaCoVacc in our study induced an immediate and potent immune responses against SARS-CoV-2 in contrast to an inactivated vaccine which required at least two or three doses of injections (Fig. 2c)^8^. We also adopted the spike pseudotyped lentivirus which encodes GFP to transduce Huh-7 cells. We found pre-incubation with 1:40 diluted sera from vaccinated mice almost completely abolished the fluorescence which was evident for placebo groups and positive controls (Fig. 2d) in line with Fig. 2c. Interestingly, sera from the vaccinated mice did not inhibit the transduction of VSV-G pseudotyped lentivirus in Huh-7 cells indicating the neutralizing antibody is spike-specific (Fig. 2d). T cell immune response is usually important for the function of vaccines to control virus infection^28^. However, in case of COVID-19, overproduction of cytokines has been correlated with the disease severity^29^. Therefore, cellular immunity must be cautious for any SARS-CoV-2 vaccines. Here, we evaluated the T cell immune response here by simulating splenocytes with a pool of spike-derived peptides. We did not found increased expression of IFN-γ and IL-2 suggesting spike-specific cellular immune response was insignificant for ShaCoVacc (Fig. 2e-f). This is line with a recent study with an inactivated SARS-CoV-2 vaccine which showed protection effects, but found no notable changes in the percentages of lymphocytes and key cytokines in vaccinated macaques^8^. Additionally, no weight loss caused by ShaCoVacc was found during the course of vaccination suggesting no evident toxicity (Supplementary Fig. 1).

**Figure 2.**
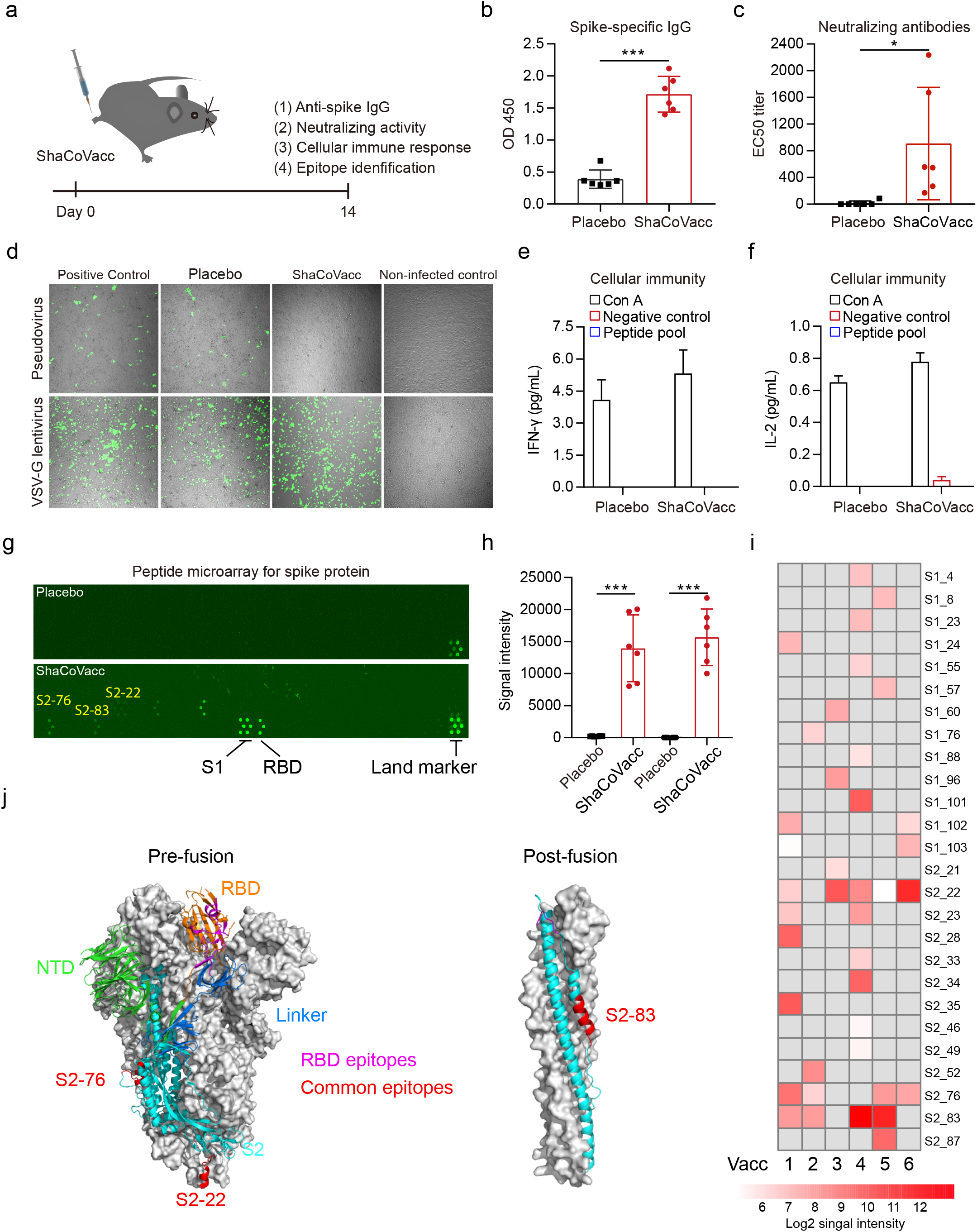
Analysis of neutralizing activity of sera from ShaCoVacc vaccinated mice and linear epitope profiling with peptide microarray. **a**, Illustration of the working plan. The sera and splenocytes were collected 14 days after footpad ShaCoVacc injection for further analysis. **b**, ELISA analysis of spike specific IgG. **c** and **d**, Neutralization activity of vaccinated sera evaluated by luciferase assay and confocal microscopy. A firefly luciferase-encoding and GFP-expressing SARS-CoV2 pseudovirus was used, respectively, to transduce Huh-7 cells. **e** and **f**, Evaluation of the spike-specific cellular immune response stimulating with a spike peptide pool. **g-i**, Peptide microarray analysis of epitopes. **g**, Representative images of spike protein peptide microarray. S1 protein and RBD were included in the microarray as controls. Highly frequent positive peptides were labeled. **h**. Antibody responses against S1 protein or RBD in vaccinated mice. Signal intensity was averaged fluorescent intensity of tinplated spots for each array. **i** Heatmap of antibody responses against peptides. Gray grid indicates negative response. **j**, Analysing the epitopes of ShaCoVacc induced spike-specific antibodies on spike protein. 6 mice used for each group, 1.5 μg ShaCoVacc or 50 μL PBS were injected via footpad into each mouse. Error bars represent ±S.D. Unpaired two-tailed student’s t-tests (**b**, **c** and **h**), *P<0.05, ***P< 0.001.

To dissect the linear epitope profile of the spike-specific antibodies in the vaccinated mice, we used a newly developed peptide microarray which contains short peptides covering the full-length of a spike. We found varying intensities of signals corresponding to certain spike peptides for the vaccinated group while no signal was observed for the placebo treated mice (Fig. 2g). We also quantified the signal intensity for antibodies against the S1 domain and receptor-binding domain (RBD), respectively. The sera from vaccinated mice elicited significantly high signals for both domains echoing our ELISA analysis of spike-specific antibodies and neutralizing assays (Fig. 2h). To access to the panorama of epitopes, we made a heat map for of all vaccinated mice and found epitope profiles for each vaccinated mouse were distinct (Fig. 2i and Supplementary Table 1). However, we also found three epitopes (S2-22, S2-76, and S2-83) that were common for 66.7% of the vaccinated mice. Interestingly, the S2-22 epitope also appeared in the majority of the convalescents uncovered by our peptide microarray (manuscript in preparation). Antibody against this epitope pulled from sera of convalescents has shown potent neutralizing activity^17^. Notably, S2-76 and S2-83 epitopes are conserved epitopes and shared by SARS-CoV and SARS-CoV-2 (Supplementary Fig. 2).

We further revealed five neutralization epitopes locating on RBD, the domain responsible for the host receptor recognition. The identified antibodies targeting these sites, therefore, can potentially block the virus entry efficiently (Fig. 2j). Notably, former studies have shown that the murine monoclonal/polyclonal antibodies that exhibit apparent binding to RBD from SARS-CoV exert no neutralization activities against SARS-CoV-2, likely caused by their distinct structural features^30^. Our findings thus pinpoint the key structural motifs in SARS-CoV-2-RBD that are susceptible to specific neutralizing antibodies. Interestingly, we also identified S2-83 epitope (N1178-V1189) locating on the HR2 region, which undergoes a dramatic refolding process during virus entry, leading to the formation of a six-helix bundle structure that finally drives membrane fusion (Fig. 2h). Considering the critical role of HR2 and its high sequence conservation among different coronaviruses (Supplementary Fig. 2), it is conceivable that antibody targeting to HR2 may potentially induce a broad neutralizing activity against various coronaviruses.

In compensation to the traditional vaccine platforms, our study provided a new vaccine platform by simulating coronavirus surface protein and internal nucleic acids, therefore, combining features of inactivated vaccines and mRNA vaccines. Due to the limited resource to SARS-CoV-2, we are currently not able to re-challenge the vaccinated animals with an authentic virus. As the VSP platform is based on lentiviral GagPol and no transfer vector is involved, it will not produce insertional and replication competent lentiviruses. Additionally, the large-scale production and quality standard of lentivirus that has been developed in CAR-T industry may facilitate the accessibility of VSP vaccine to general populations. We further revealed the epitope profiles of vaccinated mice and the epitopes susceptible to specific neutralizing antibodies, which may facilitate drug and antibody development.

## METHODS

### Cell cultures

293T and Huh-7 cells were cultured in DMEM (Gibco, USA) supplemented with 10% fetal bovine serum (Gibco, USA) and 1% penicillin/streptomycin (P/S) (Thermo Fisher Scientific, USA). Primary splenocytes and THP-1 cells were cultured in RPMI 1640 (Gibco, USA) with 10% fetal bovine serum (Gibco, USA). THP-1 cells were differentiated into macrophage-like cells with 150 nM phorbol 12-myristate 13-acetate (PMA) (Sigma) before the experiment.

### Plasmids

pCCL-PGK-spike-flag and pCCL-PGK-spike-mut-flag were constructed by replacing the GFP gene in pCCL-PGK-eGFP with spike or mutant spike (K1003P and V1004P) gene. pCMV-spike-mut-6XMS2, pCMV-spike-6XMS2-flag, pCMV-spike-mut-6XMS2-flag were generated by inserting 6XMS2 stem loop repeats between the stop codon of spike or mutant spike gene and polyA sequence while the whole expression cassette is under the control of CMV promoter.

### Production of VSP, IDLVs, and pseudovirus

VSP, IDLV, and pseudovirus were produced by 293T cells in 15-cm dishes. Cells were seeded in the 15-cm dish at a density of 10^7^/dish 24 hr before calcium phosphate transfection. The media were refreshed 12 hr after transfection. 48 hr and 72 hr post-transfection, supernatants were filtered through a 0.45-μm filter (Millipore) and ultracentrifuged at 4°C for 2 hr. Pellets were re-suspended in PBS and stored at −80°C. To produce GFP-expressing spike pseudovirus and IDLV (IDLV-spike or IDLV-spike-mut), cells were transfected with 9.07 μg pMD.2G (or corresponding spike plasmids), 7.26 μg pRSV-Rev, 31.46 μg pMDlg/pRRE-D64V, 31.46 μg pCCL-PGK-eGFP (or pCCL-PGK-spike-flag or pCCL-PGK-spike-mut-flag). To produce VSP-spike (or VSP-spike-mut), cells were transfected with 9.07 μg pMD.2G, 31.46 μg pMS2M-PH-gagpol-D64V, 31.46 μg pCMV-spike-6XMS2, or pCMV-spike-mut-6XMS2, or their flag tag versions. To produce GFP-expressing spike pseudovirus, cells were transfected with 9.07 μg pCMV-spike, 7.26 μg pRSV-Rev, 31.46 μg pMDlg/pRRE and 31.46 μg pCCL-PGK-eGFP. To produce luciferase-encoding spike pseudovirus, 293T cells were transfected with 20 μg pcDNA3.1-SARS-Cov2-spike and 20 μg pNL4-3.luc.RE.

### Western blot

To detect spike protein associated with VSP or IDLV, we use Western Blot to detect spike protein with/without treatment of PNGase F (NEB). 100 ng particles were incubated with Glycoprotein Denaturing Buffer at 98°C for 10 min. After adding GlycoBuffer 2 and NP-40 (10%), the mixtures were incubated with/without PNGase F at 37°C for 2 hr. Mixtures were then incubated with SDS loading buffer (Beyotime Biotechnology) before sample loading. To detect spike protein expressed in cells, 293T cells were lysed in RIPA 36 hr after transfected with VSP or IDLV. The lysates were incubated with SDS loading buffer supplemented with 2.5% β-Mercaptoethanol at 37°C for 30 min without boiling. The proteins were separated by SDS-polyacrylamide gel electrophoresis and transferred to the PVDF membrane. The membrane was blocked by 5% fat-free milk dissolved in TBS/0.05% Tween-20 for 1 hr then cut off according to the marker and incubated with anti-flag monoclonal antibody (Sigma) overnight at 4°C. The membranes were incubated with anti-mouse secondary antibodies (Cell Signaling Technology) for 1 hr at room temperature. Proteins were visualized by hypersensitive ECL chemiluminescence (Beyotime Biotechnology).

### Quantitative PCR

To detected spike mRNA numbers carried by VSP, total RNAs from all samples were extracted using the Viral DNA/RNA extraction kit (TaKaRa) followed by cDNA synthesization using the HiScript Q RT SuperMix for qPCR (Vazyme, China) according to the manufacturer’s protocol. RT-qPCR was performed using qPCR SYBR Green Master Mix (Vazyme) following the manufacturer’s protocol. Plasmid pLV-PGK-S-mut diluted into copies of 10^3^, 10^4^, 10^5^, 10^6^, 10^7^ per microliter were used to make a standard curve for absolute quantification. Primer sequences are as follows, forward primer: 5’-ACAGATGAGATGATCGCCCAG-3’, reverse primer: 5’-TCTGCATGGCGAAAGGGATC-3’.

### Mice

6-8 weeks old, male, specific-pathogen-free (SPF) C57BL/6 mice were inoculated with VSP, IDLV, or PBS by foot-pad injection. Animals were sacrificed by cervical dislocation under isoflurane. The animal study has complied with the guidelines of the Institutional Animal Care and Use Committee (IACUC) of the Shanghai Jiao Tong University.

### Splenocytes isolation

Spleens were removed aseptically, placed in RPM 1640 medium, gently homogenized, and passed through the cell strainer (Jet Bio-Filtration) to generate a single-cell suspension. Erythrocytes were rapidly washed and lysed by the RBC lysis buffer (Sangon Biotech), and the splenocytes were resuspended in 1mL RPMI 1640 medium.

### ELISA

HIV p24 ELISA (Beijing Biodragon Immunotechnologies) was used to measure the p24 level of the lentiviral particles according to the manufacturer. To detect SARS-CoV-2-spike specific antibodies *in vivo*, serum from the animals were used to test the spike-specific IgG by Mouse IgG ELISA (Bethyl) with a few modifications. 200 ng recombinant spike proteins were coated in 96-well ELISA plates overnight at 4°C in a carbonating buffer (pH 9.5). The plates were blocked with PBS containing 0.05% Tween 20 (PBS-T) and 2% bovine serum albumin (BSA) for 1 hr. Serum samples were diluted 1: 4 using PBS incubated for two hours, then washed and test follow the manufacturer’s instructions. To find the involvement of cellular immunity, the cytokine production in the splenic cells upon treatment of spike peptides in vitro was measured. For the IFN-γ and IL-2 assay, the splenocytes were plated in a 96-well plate at a density of 5× 10^5^ cells/200 μL. Cells were incubated with a pool of SARS-CoV-2 spike peptides of 1 μg/mL final concentrations for each peptide for 24 hr, or 5 μg/mL concanavalin A (ConA) (Sigma) and culture medium as controls. A pool of 210 peptides (15-17-mer) overlapping by 10 spanning the entire SARS-CoV-2 spike protein were synthesized and kindly provided by Dr. Shengce Tao from Shanghai Jiao Tong University, China. Peptides were pretreated with 100 μg/mL polymyxin B (Beyotime) for 30 min on ice before stimulation to exclude the LPS contamination. Cytokine IFN-γ and IL-2 in the supernatants were detected using the corresponding ELISA Kits according to the manufacturer’s instruction (MultiSciences Biotech). A standard curve was established according to the OD values, and antibody concentrations were calculated.

### Neutralization assay

To determine the serum neutralization activity against GFP-expressing spike pseudovirus. Vaccinated mouse serum (40 × dilutions) were incubated with GFP-expressing spike pseudovirus at 37°C for 1 hr before adding the mixtures to Huh-7 cells (4×10^4^ cells per well in 48-well plates). The media were changed after 12 hr and photos were token at 48 hr post infection. For luciferase-encoding spike pseudovirus neutralization assay. Serial dilutions of ShaCovacc or placebo vaccinated mouse serum were incubated with luciferase-encoding pseudovirus at 37°C for 1 hr before adding to Huh-7 cells (10^4^ cells per well in 96-well plates). The culture media were refreshed 12 hr post-infection, which was followed by an additional 48 hr incubation. Huh-7 cells were subsequently lysed with 50 μL lysis reagent (Promega), and 30 μL lysates were transferred to 96-well Costar flat-bottom luminometer plates (Corning Costar) for the detection of relative light units using the Firefly Luciferase Assay Kit (Promega) with an Ultra 384 luminometer (Tecan). A nonlinear regression analysis was performed on the resulting curves using Prism (GraphPad) to calculate half-maximal inhibitory concentration (EC50) values.

### Immunofluorescence imaging

293T cells were seed to 48-well plates with 0.1 mg/mL poly-D-lysine coated cover glasses at a destiny of 4× 10^4^/well. On the next day, cells were transduced by 150 ng IDLVs or VSPs, or transfected by 0.6 μg pCMV-spike-6XMS2 or pCMV-spike-mut-6XMS2 plasmids. Cells were fixed using 4% paraformaldehyde 36 hr after transduction and transfection. Cells were then stained by anti-flag tag antibody (Proteintech) followed by Alexa Fluor 555 IgG incubation (Cell Signaling Technology) and nuclei staining with DAPI (Beyotime Biotechnology). The imaging was performed on a confocal microscope (A1Si, Nikon) to verify the expression of Spike proteins.

### Statistics

Data are presented as mean ± S.D. in all experiments. One-way analysis of variance (ANOVA) or student’s t-tests were performed to determine the P values. *indicates statistical significance (*P< 0.05, **P< 0.01, ***P< 0.001, n.s.=non-significant).

### Data availability

Data generated or analysed during this study are available from the corresponding author on reasonable request.

## Acknowledgement

Y.C. is supported by National Natural Science Foundation of China [31971364], Pujiang Talent Project of Shanghai [GJ4150006], Shanghai Municipal Natural Science Foundation [BS4150002] and Startup funding from Shanghai Center for Systems Biomedicine, Shanghai Jiao Tong University [WF220441504]. J.H. is supported by the National Natural Science Foundation of China [81970766 and 81670818] and the Shanghai Rising-Star Program [18QA1401100]. T.Y. is supported by the National Key R&D Program of China [2019YFA0904400] and National Natural Science Foundation of China [81822027, 81630090].

## Author contribution

D.Y, S.L., and Y.C. conceived the study and designed the experiments; D.Y., S.L., X.T, Y.L., Z.X., J.H., X.Z., X.W., and J.H. performed the experiments; all the authors analysed the data; D.Y., S.L., T.Y., and Y.C. wrote the manuscript with the help from all the authors.

